# Genetic variants influence on the placenta regulatory landscape

**DOI:** 10.1101/432211

**Authors:** F. Delahaye, C. Do, Y. Kong, R. Ashkar, M. Sala, B. Tycko, R. Wapner, F. Hughes

## Abstract

**Background:** From genomic association studies, quantitative trait loci analysis, and epigenomic mapping, it is evident that significant efforts are necessary to define genetic-epigenetic interactions and understand their role in disease susceptibility and progression. For this reason, an analysis of the effects of genetic variation on gene expression and DNA methylation in human placentas at high resolution and whole-genome coverage will have multiple mechanistic and practical implications.

**Results:** By producing and analyzing DNA sequence variation (n=303), DNA methylation (n=303) and mRNA expression data (n=80) from placentas from healthy women, we investigate the regulatory landscape of the human placenta and offer analytical approaches to integrate different types of genomic data and address some potential limitations of current platforms. We distinguish two profiles of interaction between expression and DNA methylation, revealing linear or bimodal effects, reflecting differences in genomic context, transcription factor recruitment, and possibly cell subpopulations.

**Conclusions:** These findings help to clarify the interactions of genetic, epigenetic, and transcriptional regulatory mechanisms in normal human placentas. They also provide strong evidence for genotype-driven modifications of transcription and DNA methylation in normal placentas. In addition to these mechanistic implications, the data and analytical methods presented here will improve the interpretability of genome-wide and epigenome-wide association studies for human traits and diseases that involve placental functions.

**Author summary:** The placenta is a critical organ playing multiple roles including oxygen and metabolite transfer from mother to fetus, hormone production, and vascular perfusion. With this study, we aimed to deliver a placenta-specific regulatory map based on a combination of publicly available and newly generated data. To complete this reference, we obtained genotype information (n=303), DNA methylation (n=303) and expression data (n=80) for placentas from healthy women. Our analysis of methylation and expression quantitative trait loci (QTLs) and correlations between methylation and expression data were designed to identify fundamental associations between genome, transcriptome, and epigenome in this key fetal organ. The results provide high-resolution genetic and epigenetic maps specific to the placenta based on a representative ethnically diverse cohort. As interest and efforts are growing to better understand the etiology of placental disease and the impact of the environment on placental function these data will provide a reference and enhance future investigations.

## Introduction

Functional genomic approaches using combinations of genomic, epigenomic and transcriptomic data can define regulatory landscapes and provide a basis for biomarker discovery and insights into disease etiology. Advances in sequencing technologies have resulted in a surge in the number and complexity of multi-omics datasets, providing unprecedented opportunities to map the regulatory landscape, while imposing substantial analytical challenges. Increasingly, researchers acknowledge the limited usefulness of genome-wide approaches that examine a single component of transcriptional regulation in isolation and appreciate the growing importance of “multi-omic” analyses. Indeed, genomic sequence variation, epigenetic and post-translational regulators are interdependent and jointly contribute to the normal functioning or dysfunction of a tissue, calling for integrative approaches to fully appreciate the interactions between these different layers of regulation.

Originally two main lines of investigation were pursued - association studies and molecular quantitative trait loci (QTL) analysis. Association studies aim to identify traits associated with genetic variants (Genome Wide Association Studies, GWAS) and more recently traits associated with changes in DNA methylation (Epigenome Wide Association Studies, EWAS). Over the past decades, thousands of common genetic variants associated with specific diseases or phenotypes have been cataloged [1–3]. Building on the GWAS model, EWAS rapidly became a fast-growing area of research [4]. However, these discovery approaches provide limited insights into causal mechanisms. To address these limitations, QTL analyses have been implemented linking genetic variants with changes in expression (eQTL), DNA methylation (mQTL; alternatively abbreviated as meQTL) or other transcriptional regulatory mechanisms. Catalogs of thousands of tissue-specific and shared regulatory eQTL variants at high resolution, such as in the Genotype Tissue Expression (GTEx) catalog, provide insights into the diversity and regulation of gene expression across various tissues [5]. Although cytosine methylation varies with sex, age or exposure to environmental factors [6] and changes in methylation patterns have been associated with many common diseases [7], DNA methylation is also under strong genetic influences [15], and locus-specific methylation levels are often correlated in related individuals [8, 9]. Twin studies provide additional evidence of the underlying genetic effect on DNA methylation pattern [9, 10] and it has been estimated that more than 30% of the methylation variance can be attributed to genotype [11].

More directly, genetic variants and haplotypes have been shown to influence local DNA methylation patterns, most often in *cis*. Loci, where DNA methylation is under genetic control, are known as methylation quantitative trait loci (mQTL). The physical counterpart of *cis*-acting mQTLs, which can be directly scored by bisulfite sequencing (bis-seq; methyl-seq) is haplotype-dependent allele-specific methylation (hap-ASM). We (CD, BT) have previously characterized mQTLs and hap-ASM in multiple human tissue types, including T lymphocytes and brain cells, as well as a small series of placentas [12, 13]. In that work, we noted that an analysis of a larger independent series of placentas was in progress, which is represented by the current study.

Unlike most other human tissue types, the number of genome-wide –omic level studies involving placentas is relatively limited with placenta being absent from most initiatives such as the GTEx consortium or the GWAS database. However, in view of its critical role, there is growing interest in better understanding the impact of placenta malfunctions throughout fetal life. During a key developmental window, the placenta controls fetal access to nutrients, hormone production and mitigation of adverse effects from the environment, with placental dysfunction resulting in chronic diseases such as heart disease, type 2 diabetes or cancer [14–17]. Evidence suggests that such perturbations of the intrauterine environment alter the appropriate genetic programming and disrupt placental and fetal development [18–20]. Although there is strong support for changes in DNA methylation to be involved, a direct association between environmental conditions, methylation alterations, and gene expression is difficult to confirm. The unique function of the placenta is reflected through its unique transcriptome and methylome profiles. In a previous study, Storvik *et al.* [21] showed a relatively poor correlation in term of transcriptome between placenta and 34 other tissues based on microarray data with many mRNA and miRNA species specific to the placenta. Similarly, as previously shown by our group and others, the mean genome-wide level of DNA methylation across placentas is lower than that found in other tissues, with an increased representation of low and intermediate levels [13, 22]. Such singularities emphasize the need for specific investigations of the placenta regulatory landscape. Along these lines, Peng *et al.* [23] performed an eQTL study using 159 placentas. They examined correlations between GWAS data and the newly identified eQTLs and found evidence for eQTLs potentially driving postnatal disease susceptibility, supporting the Developmental Origins of Health and Disease (DOHaD) hypothesis. These findings confirm the need for a thorough characterization of the placenta regulatory landscape, preferably not limited to genetic variant and gene expression.

Among the potential limitations of mQTL and eQTL studies are subpopulation effects, cell specificity, genetic background or genomic context, as many factors that need to be accounted for to achieve reproducibility and accuracy. Further challenges include the integration of multiple types of data calls for quality control assessments, independent controls for both false-positive and false-negative findings, and a limited understanding of stochastic variation in the different signals. Finally, the availability of a reference dataset is key to support ongoing functional genomics analysis. However, the cell dependent aspect of transcriptional regulation does not allow for direct carryover of findings from one tissue or cell type to another. Lack of ethnic and racial diversity can also be a limitation, with the most comprehensive studies up to date involving predominantly Caucasian populations [24–26].

These considerations have motivated the present study where we set out to generate essential reference data sets of genotypes, gene expression, and DNA methylation patterns, based on a representative population of placentas; an understudied yet important tissue. In addition, we aimed to develop stringent analytical methods to serve as a blueprint for future studies. We intensively characterized the effects of genetic variation on gene expression (eQTLs) and DNA methylation (mQTLs) and developed a comprehensive approach to identify methylation-sensitive transcription (abbreviated here as expression-relevant Quantitative Trait Methylation; eQTM) in normal human placentas. We emphasize technical caveats, independent validations, and consideration of both false-positive and false-negative findings. Our integrative multi-omic approach provides a multilayered view of gene regulation influenced by common genetic variants in the human placentas, thus providing a basis for future advances in individualized medicine.

## Results

### Cohort information

Actual functional genomics studies are suffering from the overrepresentation of Caucasian backgrounds, which limit the reproducibility of the findings in more heterogeneous and representative populations. Aware of this limitation, it was important to characterize the ancestral history of our population to confirm the mix-population background of our cohort and to identify outliers that should be excluded from further analysis. We performed local ancestry deconvolution using PCA analysis based on the reference population using genotype data from the 1000 Genomes Project from Caucasians (CEU), Africans (YRI) and East Asians (CHB + JPT). Our data showed a clear clustering to populations, with some enrichment for mixed populations represented as expected (**S1C Fig**) validating the use of our cohort as a representative cohort for control placenta samples.

To assure the relevance of our human placenta cohort as representative of ’normal’ individuals, we used quality control measures to assess the impact of variability in the datasets. Sample heterogeneity was estimated running Pearson’s correlation across expression and methylation data. Global correlation among our samples ranges between 0.92 and 0.99 for expression values and between 0.93 and 0.99 for DNA methylation values (**S1A-B Fig**). As correlation can be artificially over-estimated when interrogating large datasets with extreme values, further analysis aimed at selecting those genes and loci with greater variability. Variability was estimated using median absolute deviation (MAD) as previously described [27]. After assigning a variability score to each interrogated CpG or transcript, variability was divided using quartile distribution creating 4 categories: no variability, low, medium and high variability. Loci and transcripts from the 4^th^ quartile of variability (high) were used to run correlation using a similar approach and range from 0.75 to 0.96 for expression values and from 0.64 to 0.98 for DNA methylation values. Principal Component Analysis (PCA) was then used to describe variability across samples and link variability to known covariates including batch effects, biological and clinical cofactors. As expected, the batch of sequencing is contributing to most of the variability in both expression and methylation datasets (**S2 Fig**). Interestingly, when focusing on the principal component accounting for most of the variability, we also found a significant association with preeclampsia status and mode of delivery in both datasets (**S2 Fig**). However, our approach still suffers from limitations as it only provides us with information on known confounders. For example, even though all samples were taken from the same region of the placentas (Methods), heterogeneity in tissue composition likely exists and could influence inter-sample variability but is not reflected in this analysis.

### Transcriptome and methylome profiles in human placenta

The placenta presents a unique transcriptome and methylome profile as demonstrated by the poor correlation in term of transcriptomic profiles between placenta and 34 other tissues based on microarray data [21]. Among these tissues, the lung was the most like placenta and liver showed the weakest correlation. Because microarrays do not query the full extent of the transcriptome, we decided to perform a similar correlation analysis using RNAseq data comparing placenta transcriptomes from our study with transcriptomes of the 53 available tissues from the GTEx consortium. For each tissue, we considered median TPM values across samples and only processed genes that were expressed (TPM>0.1) across each tissue type. The correlation scores ranged from 0.02 to 0.27 reflecting a high diversity between placenta and the other tissues (**S1 Table**). Interestingly, among the tissues overlapping between the GTEx consortium and the one tested by Storvik *et al.*, we confirmed lung as being closest to the placenta and liver to be the one with the weakest correlation. The mean genome-wide level of DNA methylation across placentas is lower than that found in other tissues, with an increased representation of low and intermediate levels [13, 22].

High variability in DNA methylation has been studied in CD34+ hematopoietic stem and progenitor cells [27] and proposed as a marker that can help localize key regulatory elements involve in cell-lineage commitment and cell specific functions. Therefore, we decided to further investigate genes with increased variability in expression or DNA methylation levels across samples and to look for associated pathways. As described above, gene and CpG were classified based on their variability across samples using MAD. We then focus on the top 500 variable genes or CpGs and performed pathways enrichment analysis (**S2, S3 Tables**). Traditional gene set enrichment analysis (GSEA) does not take into account the physical characteristics of the gene and has been shown to be biased by factors such as the length of the gene [28]. To address this, we used the Bioconductor package GOseq [29] developed to control for variability of the length of genes. For each CpG, the closest associated gene was considered. For the CpG analysis, we adapted the original version of GOseq to control for the number of probes by gene instead of gene length, as length is not relevant in this case. Interestingly, in both datasets, the majority of the pathways (**S4 Table**) were related to the immune response involving genes from the HLA family suggesting that heterogeneity among samples could be driven by a different level of inflammations in each placenta. Looking only at expression data, the top pathway is KEGG “ECM-receptor interaction”. ECM related genes have been previously associated with placental development [30] and have been shown to be correlated with oxygen tension and nutrient availability [31]. Therefore, variability in the expression for these pathway-associated genes may reflect various *in utero* exposures with impact on placenta development and potential long term consequences even in the absence of clear phenotype at birth.

### Expression and methylation quantitative trait loci in human placenta tissue

To generate maps of associations between genetic variants and gene expression or CpG methylation patterns specific to the human placenta, we performed eQTL and mQTL analysis across 1,374,581 SNPs, 23,003 genes, and 485,578 CpG sites respectively. We focused on identifying cis-eQTLs within windows of 1 Mb between the gene transcription start site (TSS) and each SNP. QTL tools identified 28,906 significant associations with 1,916 preserved after permutations, which was further reduced to 985 eQTLs after exclusion of associations that failed the quality threshold (**Methods and** **Fig 1**, **S3 Fig**). A complete list is of the 985 eQTLs used for further analysis is provided in **Supplementary Table 5.** For mQTL analysis, we focused on 150 kb windows (75 kb upstream and downstream of the CpG site) as it has been previously published that mQTLs correlate best with SNPs within 1-2 kb and that most mQTLs are located within 100 kb in several tissues including placenta [13, 32, 33]. 471,852 significant associations were called using our pipeline, only 105,031 associations pass the permutation test with 4,342 mQTLs conserved for further analysis (**Fig 1**, **S3 Fig**) after stringent exclusions based on quality threshold (**see Methods**) and potential probe artifacts. The 4,342 mQTL associations kept for further analysis are recapitulated in **Supplementary Table 6**.

**Fig 1.**
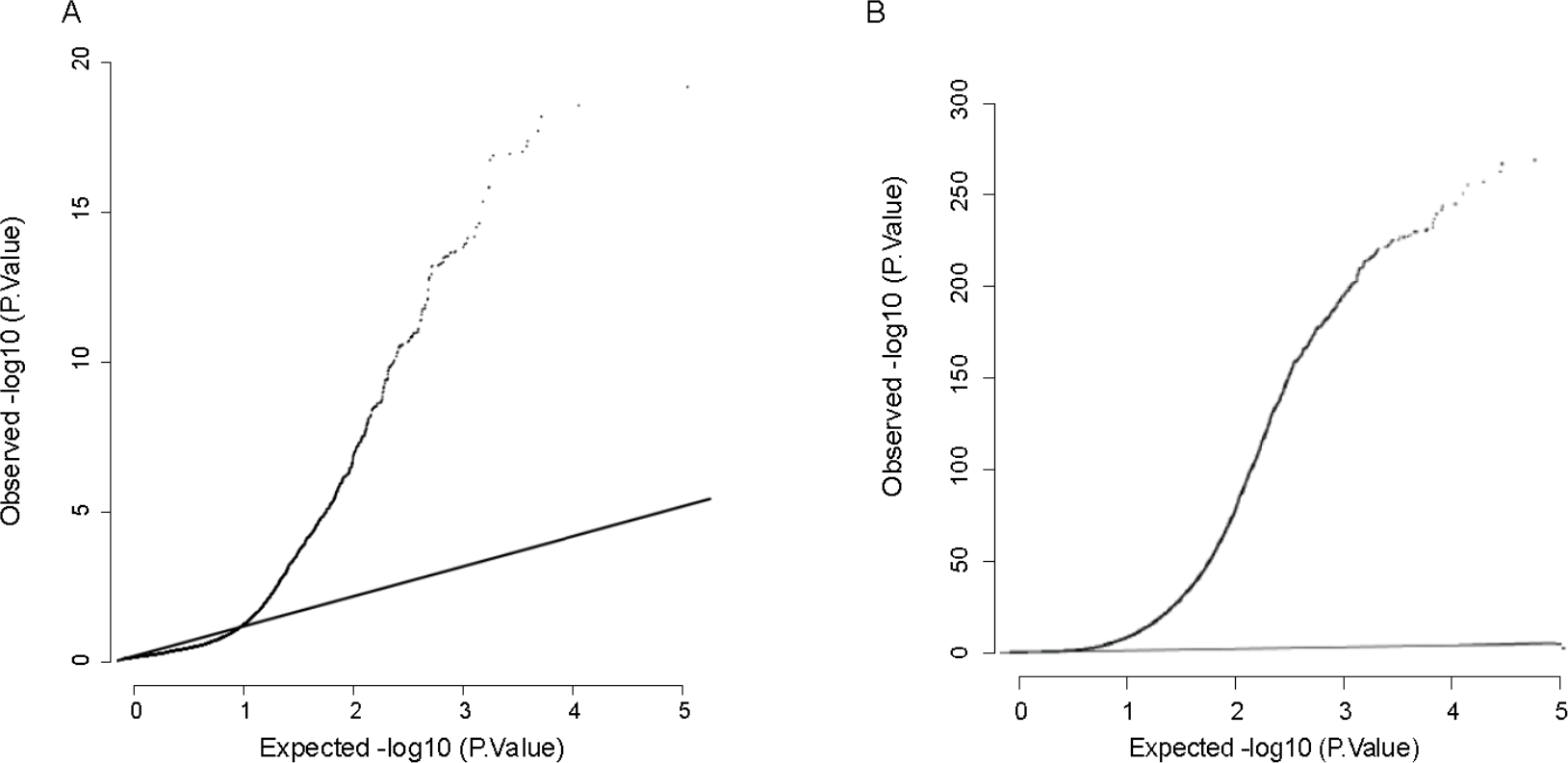
Identification of eQTLs and mQTLs. Q-Q plot representing expected versus observed significance for the association between gene expression and genetic variant (A) and between CpG methylation and genetic variant (B)

Some genetic variants that appear as individual differences in methylation can act trivially via SNPs that create or abolish CpGs (CpG-SNPs) [34] or via technical artifacts due to incomplete probe binding [34] or probe-cross reactivity [35] referred to here as “at risk-probe”. Incomplete probe binding may suggest the presence of a genetic variant within the probe binding site with greater impact in the proximity of the 3’ end of the binding probe with differences in signal intensity persisting for up to an approximately 20 bp [36]. Therefore, to make our lists more stringent, we identified and excluded associations involving probes that contain known SNPs, along with probes previously identified as cross-reactive. A list of excluded probes is in **Supplementary Table 7**. As we illustrate by independent validations below, this high level of stringency is expected to reduce false-positives. Additionally, it can simultaneously give rise to false-negative findings, particularly for loci queried by probes that contain internal SNPs with modest minor allele frequencies and/or those without strong effects on probe binding, which can yield accurate readouts in most samples.

By using a reference-based approach, identification of probes sensitive to CpG-SNPs may be subject to tissue and sample specificity. Recently, a methodology has emerged to identify “at risk” probes independently of reference datasets, based on a clustered distribution of methylation scores [36]. Using this approach, the authors were able to identify a significant number of “gap probes” that largely overlap with CpG-SNP probes. However, as SNP-affected probes can exist without gap signatures, the implementation of this reference free approach should be considered in complement with the reference-based approach. To be as inclusive as possible, we applied the “gap-hunter” [36] approach to our datasets to identify remaining probes and associations subject to possible technical artifacts. By doing so, we were able to detect 10,971 “gap probes”. From these, 3,235 are overlapping with previously identified CpG-SNPs probes and 205 with cross-reactive probes. Finally, 432 of the remaining “gap probes” overlap with our 4,342 mQTL associations and 158 overlap with our eQTM associations. These probes were not excluded from further analysis but were annotated following the recommendation of Andrews *et al*. [36] (S6, **S7 Tables**).

Interestingly, after FDR correction, about 22% of the mQTL (23,043 out of 105,031) involved an association between an at-risk probe and genetic variant. We are aware that this proportion reflects in part the stringency of our criteria for exclusion. However, these findings are strong enough to encourage for stringent criteria when running genome-wide analysis. To take into consideration false-negative results, it is particularly attractive to use annotation instead of direct exclusion, as we have used previously [13], and in part adopted here.

### Reproducibility of QTL associations

To assess the robustness of our approach we were first interested in looking into the overlap between our newly identified associations and published data on the human placenta. Because identification of the causal genetic variant is still challenging due in part to linkage disequilibrium, overlaps based on genetic variants are likely to not be as informative. Therefore, we decided to look at overlap driven by gene or CpG for eQTL and mQTL associations respectively. A recent study published by Peng *et al.* [23] has identified 3,218 eQTLs associations based on 159 human placenta tissues. These data represent a unique opportunity to assess data reproducibility by examining the number of associations preserved across different cohorts. Our 985 eQTL associations represent 615 unique genes and 62% of them overlap with genes previously identified by Peng *et al*. (**S8Table**). As noted above, we (CD, BT) previously reported a list of mQTLs in placentas from a smaller group of samples [13], independent of the current series. Using more lenient criteria for probe exclusions, we identified 866 mQTLs (n=665 with similar exclusion criteria). Among these mQTLs, 319 (~48%) were also found in the current series, with many of the non-overlapping “hits” reflecting probes on the 450K methylation arrays that were excluded from the current study (**S9 Table**). The significance of these overlaps was further confirmed using a permutation test (n=1,000). The significance of the enrichment was defined by the overlap between observed versus expected distribution. Here, the null distribution represents a random sampling from the total number of annotated Refseq genes or CpGs interrogated by the Illumina 450K platform. Random permutations failed to replicate such overlap with a maximum of 17% overlapping genes and 2% overlapping CpGs. Therefore, we feel confident in the strength of our study and believe that these newly identified associations represent a solid foundation for further exploration.

After demonstrating a significant overlap for placenta in different cohorts, we were interested in evaluating the overlap across tissue types. The GTEx Project, assessing and cataloging eQTLs across more than 40 tissues, is a valuable resource for studying human gene expression regulation and its relationship to genetic variation across tissue types. However, among the list of tissues in the GTEx project, the placenta has so far been missing. Therefore, our current study represents a key opportunity to expand the GTEx exploratory work toward this crucial fetal organ. To evaluate to what extent the placenta eQTLs replicated the significant SNP-gene pair eQTLs found in the GTEx study (at FDR<5%) we used the π1 statistic. The π1 statistic can be interpreted as the fraction of eQTL associations shared between the placenta and the other tissues available through GTEx. This approach allowed us to identify placenta-specific eQTLs, and identify tissues with similarity to the placenta in this respect. Levels of significance were collected across the different tissues for our list of significant eQTL association from the placenta. π1 value range from 0.31 for the brain cerebellar hemisphere tissue to 0.69 for the transformed fibroblast cells (**S4 Fig, S10 Table**). It is interesting to note the absence of interdependence between gene expression correlation and number of conserved eQTLs. Indeed, no significant overlap was found (Fisher’s exact test, p.value=0.5935) when overlapping the most similar tissues to the placenta (n=10) based on π1 value (**S10 Table**) with the most similar tissues to the placenta (n=10) based on gene expression correlation value (**S1 Table**). Among the 985 eQTL associations identified in placenta, 63 appear to be specific to placenta as not found significant in any other GTEx tissues (**S11 Table**). Unfortunately, similar curated reference doesn’t exist for mQTL associations with only a limited number of tissue types studied (brain, blood, lymphoblastoid cells, T-cells and lung [37–42]). Contrary of the GTEx analytical design, there was no consensus on how associations were defined increasing variability across studies, a likely confounder when assessing the biological relevance of the overlap.

### Evaluation of stringency thresholds by bisulfite sequencing

In this study, we used a conservative approach to minimize false positive by excluding probes with potential cross-reactivity or mapping of common SNPs within 20 bp of the queried CpG, which might affect the probe hybridization. This stringent approach will reduce false-positives, but it can also lead to false-negative findings in which *bona fide* mQTLs are discarded. To investigate this aspect directly, we applied targeted bisulfite sequencing (bis-seq) to assess allele-specific CpG methylation (ASM), which is the physical counterpart of mQTLs. For this purpose, we chose three loci that passed our current stringent criteria for inclusion of the 450K probes and showed mQTLs (ranking from 82 to 1900 in strength, see S6 and S7 Tables), and four loci for which the probes were excluded by the current criteria but were included by our more lenient criteria and showed mQTLs in our prior study of the smaller placental series [13]. While the net methylation at the index CpGs showed more inter-individual variability in the bis-seq data, the average net methylation matched that observed in 450K arrays (+/− 5%) suggesting no or mild hybridization issues with the four excluded Illumina probes. Further, statistically significant ASM, affecting from 22% to 45% of the heterozygous samples, was found at the amplicon level for all 7 loci (**S12 Table and** **Fig 2** **and S5, S6 Figs**), thus validating these loci as genuine mQTLs.

**Fig 2.**
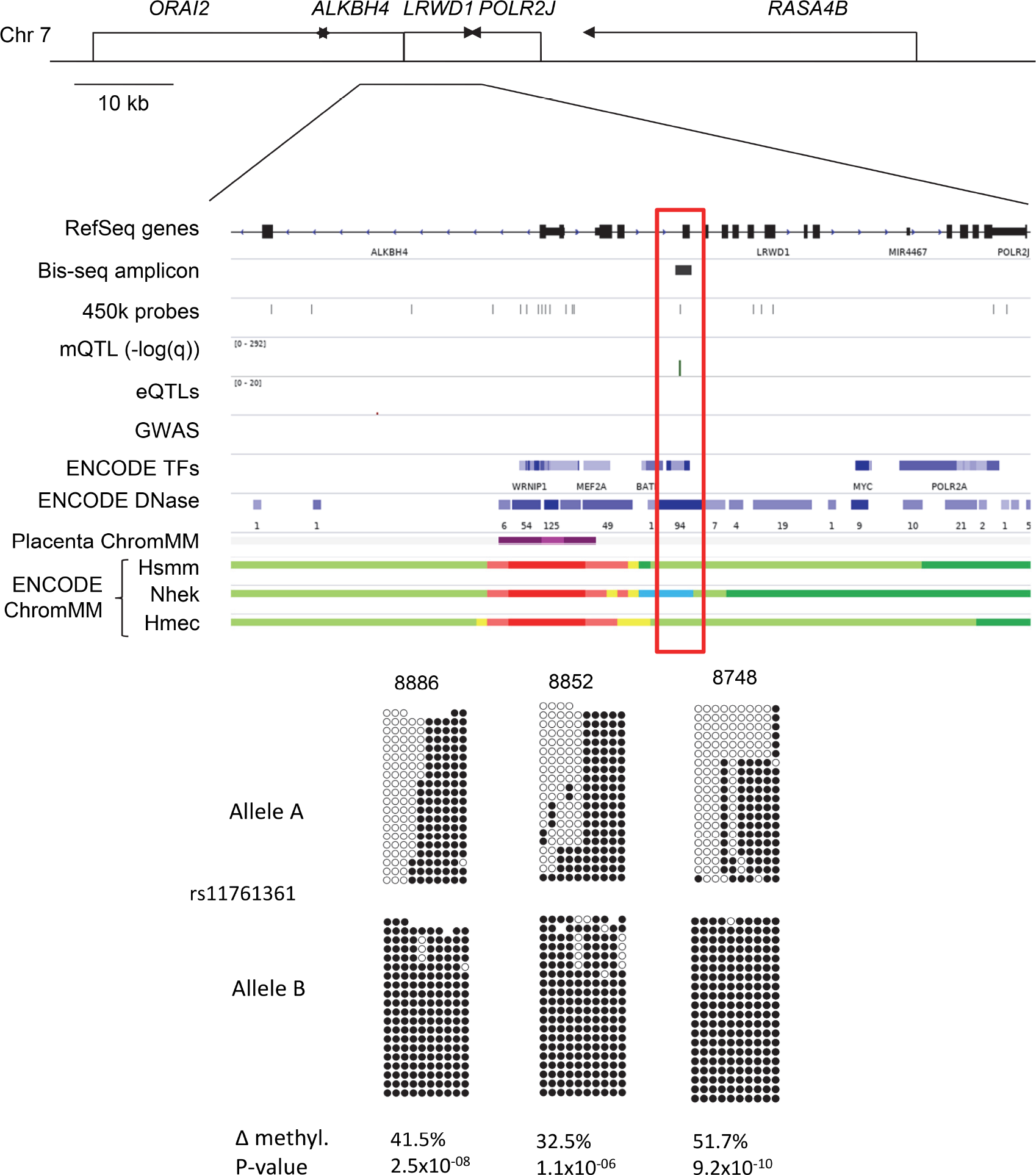
mQTL validation. Targeted bis-seq data showing Hap-ASM in *LRWD1* region. The bis-seq amplicon covers 289 bp including the mQTL index CpG queried in Illumina 450K BeadChips arrays (cg25932869), as well as contiguous CpGs. Graphical representation from 3 representative heterozygous placenta samples with hap-ASM is shown. This region overlaps with the common SNP, rs11761361. Allele A and B are analyzed and represented separately. The SNP dictates methylation level with the alternate allele (allele B) being significantly hypermethylated compared to the reference allele (allele A), suggesting the presence of haplotype-dependent ASM in 19 out of 21 heterozygous samples. The low methylated allele is significantly biased toward allele A (p=10^−06^, using binomial test) which ruled out imprinting. The hap-ASM overlaps with an insulator chromatin state (blue) in a NHEK cell line which suggests the presence of a dynamic regulatory element. For each heterozygous sample, Wilcoxon p value and methylation difference between alleles were calculated by bootstrapping (1,000 sampling of 20 reads per allele) and are indicated only for significant hap-ASM defined as the difference in percentage methylation >20%, >3 ASM CpGs, and p < 0.05. One representative random sample of each allele (20 reads per allele) is shown. mQTL, eQTL, and placenta chromatin states tracks are from our analyses. The GWAS SNPs track was downloaded from the NHGRI-EBI GWAS Catalog [1], TF ChIP-seq data, DNAse I hypersensitive sites and cell line chromatin states were downloaded from the ENCODE project [78]. ∆Meth (difference in the percentage of methylation between alleles in heterozygous samples) and Wilcoxon p-values are from bootstrapping.

These data indicate that our stringent criteria for excluding Illumina probes, while minimizing false-positives, can also lead to false-negative findings from exclusion of probes that are, in fact, legitimately informative. Based on these findings, we conclude that an optimal approach for harvesting and interpreting mQTLs using Illumina arrays is to annotate each probe as meeting either stringent or lenient criteria for predicted reliability, and include this information as a descriptor in comprehensive lists of mQTLs. Here we adopt this approach for reporting mQTLs (**S6, S13 Tables**), but for our bioinformatic enrichment and pathway analyses, we have used only the mQTLs that pass our stringent probe selection criteria.

### Characterization of gene expression and DNA methylation patterns associated with genetic variants

Characterizing gene expression and DNA methylation patterns that are influenced by genetic variants can potentially provide functional information about disease-causing genes and pathways. Therefore, we were interested in better characterizing genes (n=615) and CpG sites (n=4,324) associated with eQTLs and mQTLs. First, we aimed to assess correlations between level of expression or level of DNA methylation with genetic variants. Genes were first assigned to different categories of expression from not expressed to highly expressed using quartiles based on our placental RNA-seq dataset. A similar approach was applied for DNA methylation levels. Enrichments for specific quartiles of expression or DNA methylation were assessed by permutation as described in Methods. The significance of the enrichment was defined by the overlap between observed versus expected distribution. In these cases, null distribution represents a random sampling from the total number of annotated Refseq genes or CpGs interrogated by the Illumina 450K platform. eQTL associations were found significantly enriched for genes with intermediate and high level of expression (**Fig 3**). Such enrichment has been previously reported in other tissues than placenta by the GTEx consortium [5]. By comparing the profile of expression between genes associated and genes non-associated to eQTLs, they showed a shift toward high expression for associated genes across tissue types. mQTL associations were enriched for CpGs with intermediate levels of DNA methylation (**Fig 3**). This pattern is likely to reflect on the placenta specific methylation profile with enrichment for low and intermediate DNA methylation levels. This can also explain the limited range of variation in DNA methylation across the different genotypes found in our selected mQTLs (with a regression slope <5% for 87% of the association). Due to the nature of the DNA methylation signal, intermediate levels of methylation are likely to reflect cellular heterogeneity [27]. Therefore, it is possible that mQTLs associations help to identify genetic variant impacting cell differentiation mechanisms, expanding downstream functional consequences associated with mQTLs.

**Fig 3.**
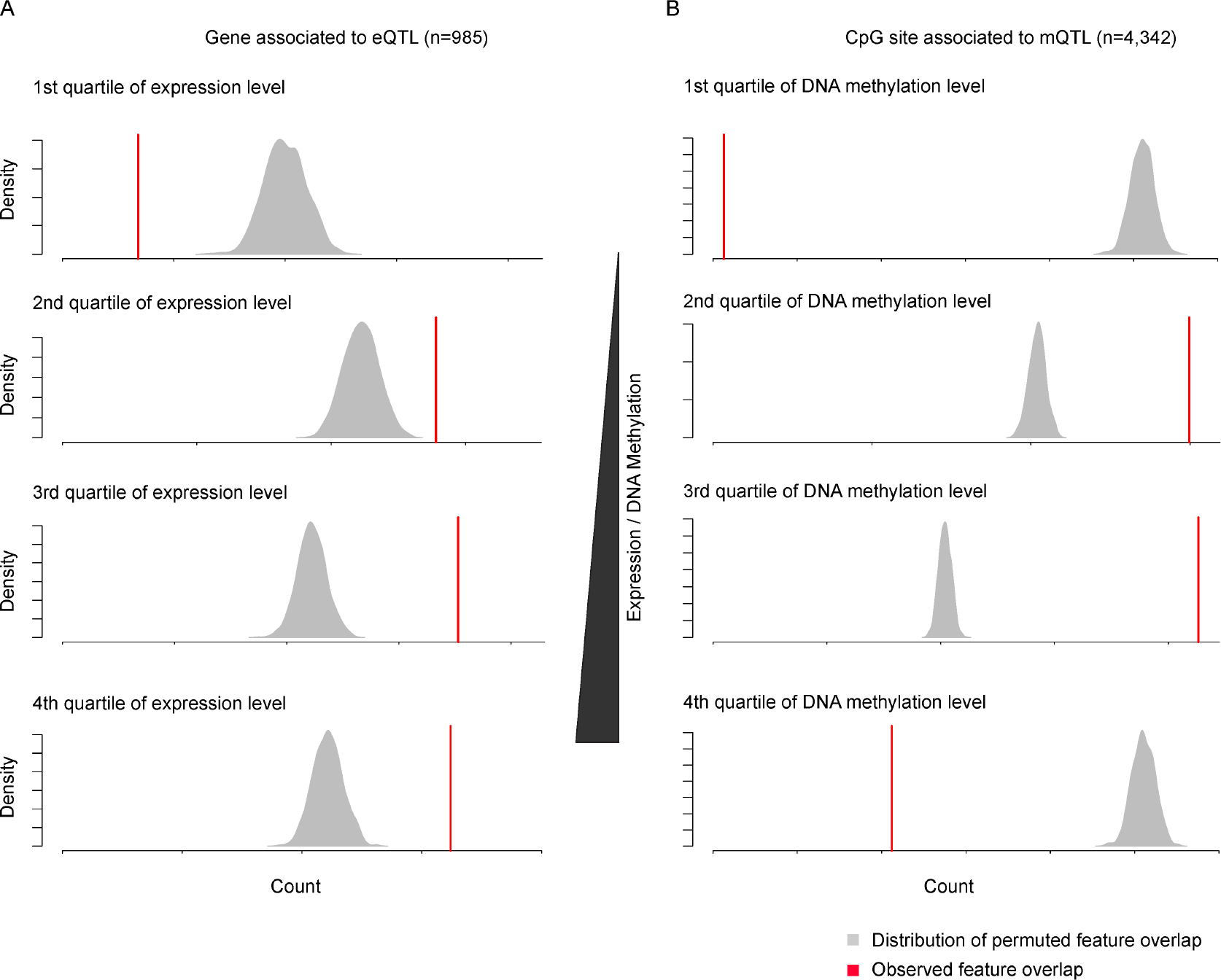
Expression and Methylation profiles associated with eQTLs and mQTLs. Enrichment for quartile of expression (A) or DNA methylation (B) was assessed using permutation tests. The density plots represent the distribution of overlaps between random sampling and the different quartiles whereas the red line illustrates the overlap value between candidate associations and corresponding quartiles.

To lay a groundwork for future studies on disease susceptibility, identification of biological pathways associated with eQTLs and mQTLs are of interest. In this context, it is valuable to identify pathways involving genes with greater sensitivity to genetic background. Therefore, we looked at gene set enrichment for genes associated with an eQTL and the nearest gene from CpG sites associated with a mQTL using the Bioconductor package GOseq [29]. For both eQTLs and mQTLs, we found enrichment for inflammation related pathways (**S14 Table**) suggesting that the variability captured by our associations may, in fact, reflect different degrees of inflammatory response across our collected samples. It is also interesting to note that the enrichment analysis outcomes (**S14 Table**) rely mainly on genes part of the major histocompatibility complex (HLA-DRB1, HLA-DBR5, HLA-DQA1, HLA-DQB1, HLA-C). These genes are known to be highly polymorphic with 13,840 different HLA alleles reported in the IMGT/HLA database [43] suggesting polymorphism mechanism as a possible bias in QTL and GWAS analysis. Finally, we decided to look into the overlap between the gene associated with eQTLs and gene associated with mQTLs where CpG was associated to the closest gene. We found 111 genes overlapping between the 2 associations (**S15 Table**) with pathway enrichment only relying on HLA related genes.

### Genetic variants associated with eQTLs and mQTLs

Genomic context is known to play a cell type-specific role in regulating transcription. To determine whether genetic variation was occurring at regulatory sites, we took advantage of public chromatin mapping data for placenta generated by the Roadmap Epigenomics Program (**S16 Table**). The DNase hypersensitivity and ChIP-seq data create combinatorial patterns that can help to define functional elements in the genome (**S7 Fig**) [44]. However, because placenta specific reference datasets are limited and do not include data from term placenta, we were compelled to build our genomic annotation based on data from the pre-term placenta which may affect the accuracy of our annotation especially in a highly dynamic tissue like placenta. For relative enrichment studies, the observed overlap between genetic variants and a genomic feature was compared with the expected overlap given the total coverage of the annotation. For permutation tests, we compared the observed overlap of SNPs significantly associated with eQTLs (n=608) or mQTLs (n=3,022) and a particular genomic interval (for example gene promoters, CpG islands) or feature from ChromHMM analysis with the distribution of the same overlap under the null hypothesis.

SNPs associated with eQTLs and mQTLs were enriched in candidate promoters (Feature 1), candidate active and poised enhancers (Feature 2-4) and enriched at regions likely to be transcribed (Feature 7) (**Fig 4A-B**). When looking at the distribution of the distance between SNP and gene for eQTL association (n=985) or SNP and CpG for mQTL association (n=4,342), we found enrichment for closest interactions (within 10Kb, **S8 Fig**) suggesting that the associated genetic variant has an effect via its presence in proximal cis-regulatory elements. We then explored the genomic context for CpGs (n=4,342) associated with mQTLs and found, as in our previous study of multiple human tissues [13], that they are enriched in candidate active and poised enhancers (Feature 2 and 3) (**Fig 3C**). This finding suggests a possible cis-regulatory impact of mQTLs on gene expression.

**Fig 4.**
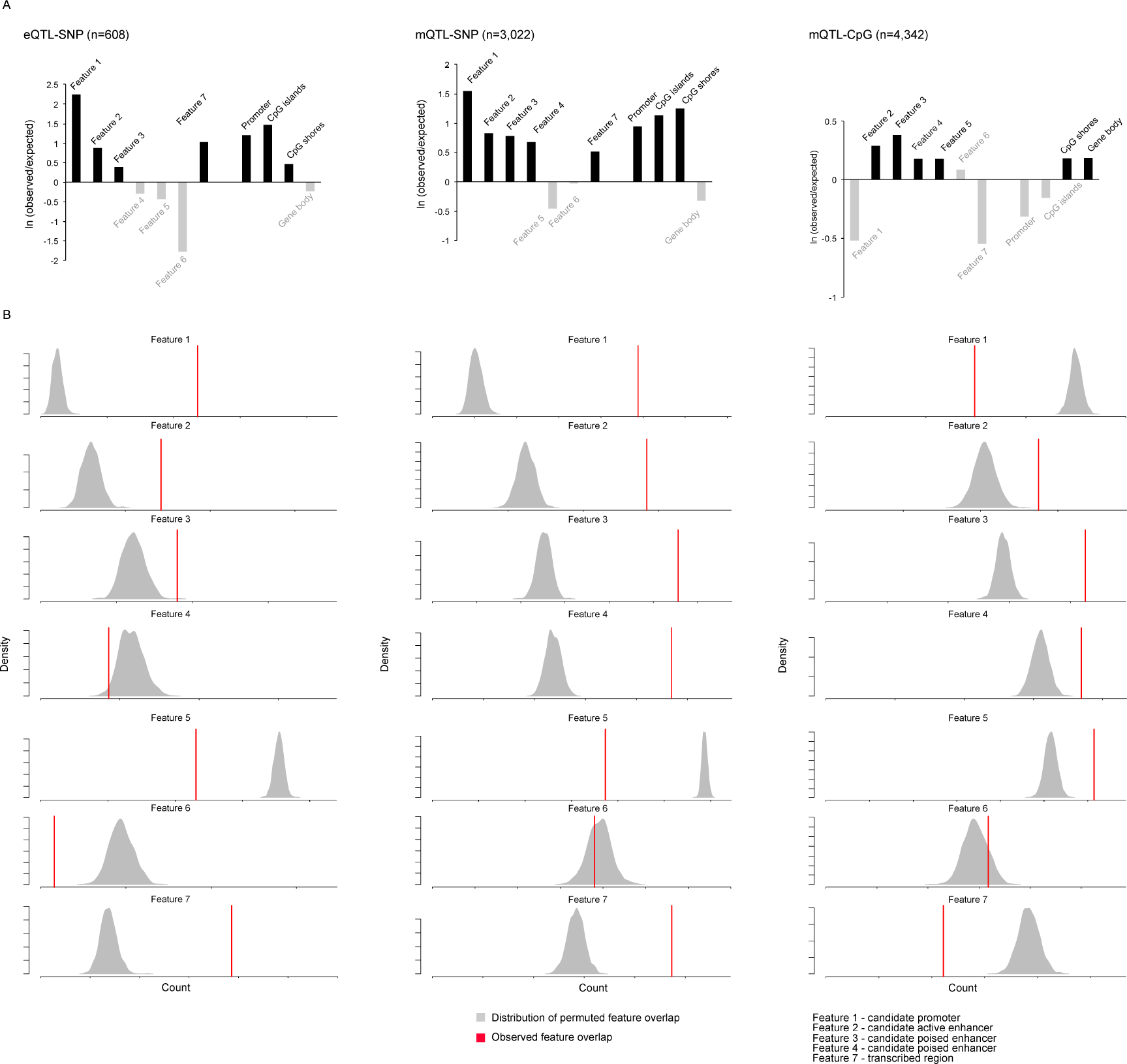
Genomic annotations of placental eQTLs and mQTLs. Bar graphs representing the observed versus expected ratio for each feature define by ChromHMM analysis and from Refseq annotations. Grey filling represents enrichment that did not reach significance (A). Significance was assessed using a permutation test. The density plots represent the distribution of overlaps between random sampling and the different features whereas the red line illustrates the overlap value between candidate associations and the corresponding features (B). eQTL-SNP refers to enrichment considering the genetic variant associated to eQTL, mQTL-SNP refers to enrichment considering the genetic variant associated to the mQTL and mQTL-CpG refers to enrichment considering the CpG associated to the mQTL.

### Transcription factor binding sites enrichment analysis

Using combined genetic and epigenetic profiling applied to other human tissues, we and others have found evidence that some of the epigenetic effects of genetic variation are mediated by SNPs in insulator elements and transcription factor binding sites [13, 45–47]. Therefore, we decided to analyze the impact of genetic variation on the integrity of the binding motif in correlation with predicted binding affinities. Due to linkage disequilibrium, the index SNP may not be causal, so we decided to also investigate the impact of previously identified SNPs in proximity to the eQTL and mQTL associated gene TSS or CpG respectively (regardless of their linkage disequilibrium status). This approach allows us to expand our discovery to transcription factors likely to be affected by linked genetic variants. We compared the predicted binding affinities between the reference and alternative alleles within a window of 39 bp around each targeted genetic variant using the motif-based sequence analysis tools FIMO from the MEME suite [48]. To assure the accessibility of the binding site, only those overlapping with DNAse hypersensitive regions (ENCODE data, release 3) were considered for further analysis. The analysis was performed at the motif level. The list of altered binding sites was further pruned down based on 2 criteria (1) the significance of the binding with the reference allele (q.value<0.05) and (2) the difference in binding affinity between the reference and alternative allele. Distributions of the difference of binding affinity between the reference and alternative sequence were used to define the cutoff for predicted allele-specific binding (difference in binding >8, **S9 Fig**). The significance of the alteration from our filter list of genetic variants and transcription factor binding motif was assessed using atSNP [49]. atSNP performs computations for between-allele score differences [50] and outputs p.values for each targeted SNP and selected transcription factor binding motif. The complete list of affected transcription factor binding sites is in **supplementary table 17**.

Interestingly, when looking at transcription factor binding sites associated with mQTLs a significant number have been previously associated with alterations of the epigenetic landscape (CTCF, REST, HDAC2, EWSR1, FLI1, SP4/1 and BHLHE40). CTCF (CCCTC-binding factor) previously identified as associated with mQTLs and haplotype-dependent allele-specific methylation (hap-ASM) by Do *et al.* [13] is one of the best documented transcription factors affecting DNA methylation. CTCF, which acts to anchor high-order chromatin loops and binds at some of its recognition sites in a CpG methylation-dependent manner, is known to have an essential role in imprinting control achieving allele-specific gene regulation [51] and CTCF binding locally influences DNA methylation [52]. REST (REI-silencing transcription factor) inactivation has been shown using a transgenic approach to induce *de novo* methylation and conversely, the expression of REST was sufficient to restore the unmethylated state of the DNA sequence bound [52]. Direct association between HDAC2 (Histone deacetylase 2), EWSR1 (Ewing sarcoma breakpoint region1), FLI1, SP4/1 and BHLHE40 (Basic helix-loop-helix family member E40) and changes in DNA methylation have not yet been demonstrated but they have been previously associated with changes in histone methylation or acetylation [53–57]. Numerous evidence supports a complex interplay between histone modification and DNA methylation (for review [58, 59]). However, if in numerous contexts a bidirectional association between changes in DNA methylation and histone modifications has been reported, the exact mechanism involved still needs to be identified. Interestingly, these associations seemed to not only rely on direct interaction between DNMT enzymes and histone methyltransferases but also on the recruitment of a variety of intermediate factors such as member of the SET domain family (G9a) or member of the Polycomb complex (PRC1) suggesting that a large number of histone modification factors could indirectly affect DNA methylation. Among the different transcription factor binding sites linked to eQTLs, we also found EWSR1, FLI1, and SP4/1, suggesting that the effect of transcription factors on gene expression may be in part mediated via epigenetic modifications. This list of transcription factors provides candidate mechanisms to explain the association between genetic variants and gene expression or DNA methylation.

### Genetic variants linked to eQTLs or mQTLs and association with GWAS peaks

The GWAS approach identifies disease-related enrichment for specific genetic variants. This approach has considerably improved marker discovery but, because of linkage disequilibrium among many SNPs in each chromosomal region, it has a limited ability to pinpoint disease genes and regulatory sequence variants. “Post-GWAS” approaches that integrate eQTLs and mQTLs to GWAS datasets can help to overcome these challenges [12]. For our current analysis, we overlapped the genetic variants listed in the GWAS catalog [60] with those associated with eQTLs (n=608) and mQTLs (n=3,022). Only 7 genetic variants associated with the placental eQTLs were previously found in GWAS peaks. Of these, three validated the previously reported association between the genetic variant and gene in the GWAS database (**S18 Table**). Two mQTLs shared a genetic variant with statistical peaks in the GWAS database (**S19 Table**). The GWAS database references a limited number of studies involving trait directly associated with placenta function or dysfunction. We identified only 2 studies related to preeclampsia and 6 related to birth weight and none of them appears in the overlaps with eQTLs and mQTLs. Nonetheless, among these hits, some are found associated with traits that have been previously linked to fetal growth, like obesity and type II diabetes where the placenta is likely to play a role. rs35694355 overlaps with our eQTLs and have been previously associated with obesity. Interestingly, this genetic variant is associated with CRACR2B gene in the GWAS database but with IRF7 in our eQTL study. Contrary to CRACR2B, evidence for a direct association between IRF7 and obesity have been shown [61] with IRF7 knockout mouse being protected from gain weight after high-fat diet exposure. IRF7 is also highly expressed in placenta and it though to play a role in the immune barrier between the fetus and the mother [62]. Due to the tight correlation between inflammatory response and obesity, IRF7 may appear as a better candidate to link rs35694355 with disease sensitivity. From our mQTL analysis, rs623323 has been previously associated with Type 2 diabetes in the GWAS database. In both datasets, rs623323 was associated with NXN gene. Interestingly in a recent publication, increase of DNA methylation and decrease of expression for NXN has been characterized in placenta from women with diabetes during pregnancy [63] suggesting that the presence of rs623323 may influence susceptibility to type II diabetes via alteration of NXN function thus providing with a great example of how integrative QTL analysis can provide with evidence to better understand disease susceptibility.

### Relationship of DNA methylation to gene expression

Being able to establish transcript-CpG specific associations (eQTMs) based on gene expression and CpG DNA methylation at a genome-wide scale in a tissue specific manner will significantly improve further interpretations of epigenetic modifications when considering their functional implications. Using our transcriptomic and epigenomic data, we decided to identify CpG methylation and gene expression with a significant correlation. To do so, we applied a similar approach to the one used for eQTL analysis. Looking at different combinations of CpG sites and transcripts with linear regression in a 100 kb window (matrixEQTL), we were able to identify 55,492 eQTM associations, representing 47,140 unique CpG sites and 14,352 unique genes. After permutation and quality control, 2,655 eQTM associations remained with 2,538 unique CpG sites and 1,269 genes (**S20 Table**). 515 associations were located in the sex chromosomes. To make the analysis less computationally intensive, we decided to sample gene expression profiles instead of CpG methylation profiles during permutation.

A genomic context-dependent correlation between DNA methylation and expression has previously been shown [64]. Negative correlation, when an increase in DNA methylation is associated with a decrease in expression, was found for CpG sites located in the promoter region. A positive correlation, when an increase in DNA methylation is associated with an increase of expression, was found for CpG sites located in the gene body. We first assessed if such profiles were also found among our associations. Both negative and positive associations were enriched for proximal interaction (**Fig 5A**). We then examined enrichment between our newly annotated features specific to placenta and our eQTM candidates stratified based on positive (n=1,009) and negative (n=1,646) correlations. We found the negative association to be enriched for feature 1 (candidate promoter) and 2 (candidate active enhancer) and associations with positive correlation to be enriched for feature 5 (**Fig 5B**). Unfortunately, feature 5 does not show a clear significant enrichment pattern for the interrogated histone marks (**S7 Fig**). However, looking at the distribution over Refseq annotations, we confirm the enrichment for gene body region. This validates the inversely correlated relationship between DNA methylation and gene expression in promoter regions as well as the positive correlation when targeting the gene body region.

**Fig 5.**
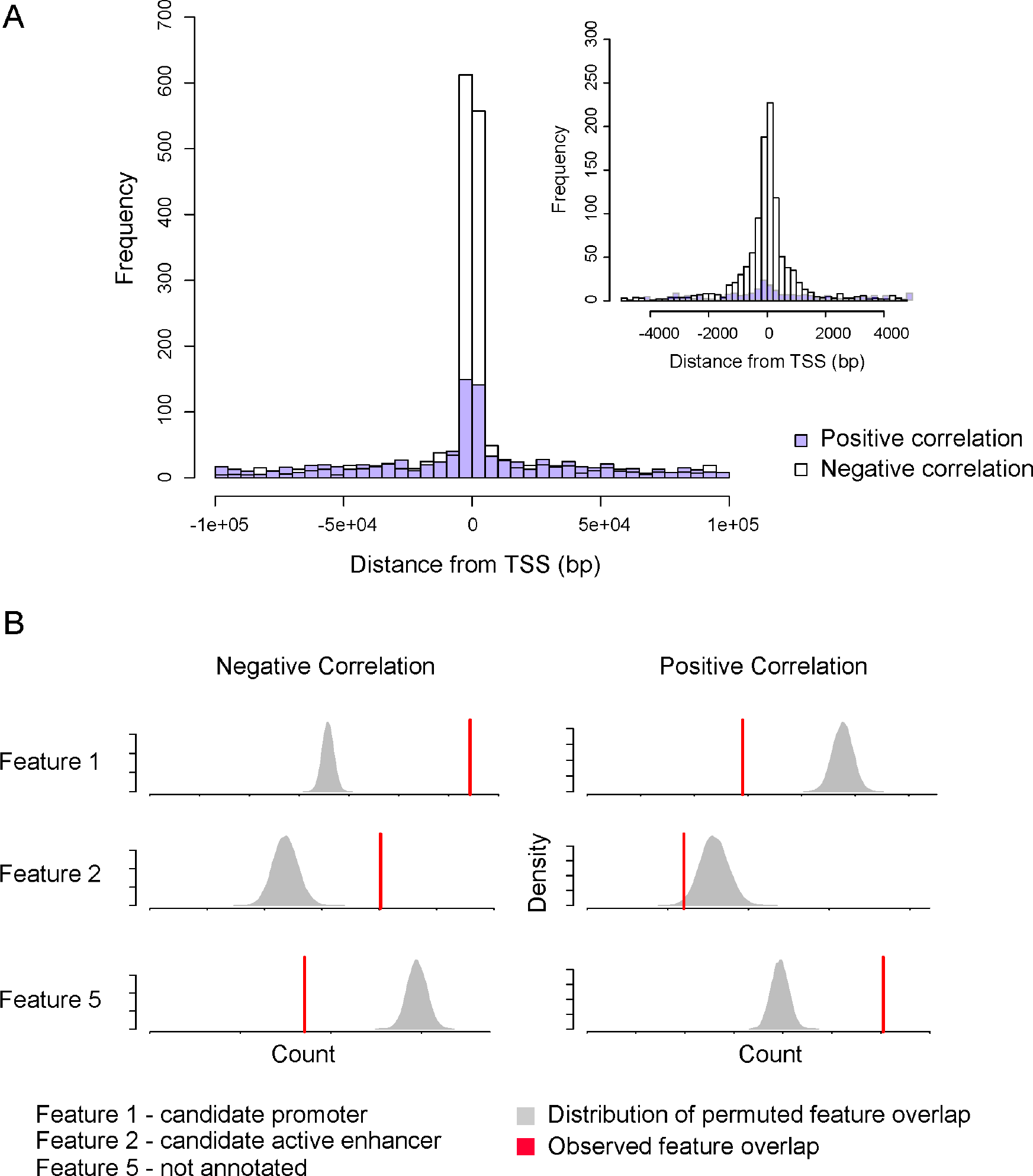
Placenta-specific genomic annotations of eQTMs. The histogram represents the density of eQTM associations in function of the distance from transcription start site (TSS) considering positive and negative correlation separately (A). The density plots represent the distribution of overlaps between random sampling and the different features as defined by permutations tests whereas the red line illustrates the overlap value between candidate associations and corresponding features (B).

To identify potential mechanisms underlying the correlation between DNA methylation and expression, we assessed the role of transcription factors, looking at enrichment for specific binding sites overlapping the CpG site from the eQTM associations (n=2,655) using a similar approach as described above. The null distribution represents the sampling of CpGs (n=1,000) from the population of all assayed CpGs overlapping with a specific TFBS. Associations with negative correlation were found to be globally enriched for transcription factor binding sites suggesting a general mechanism where increase DNA methylation at a given binding site will alter binding abilities and affect transcription. Interestingly, when looking at associations that showed a positive correlation between expression and DNA methylation, we found enrichment for only one transcription factor, ZNF217 (**S21 Table**). The ZNF217 transcription factor has been previously recognized as a human oncogene and is known to be part of a complex that contains several histone-modifying enzymes strongly associated with gene repression [13]. Thus, alteration of the binding of this transcription factor is predicted to disturb the formation of a gene repression complex and lead to an increase of expression for the targeted gene by diminishing its inhibition. There is also evidence for a more distal action of ZNF217 with only 2% of the binding site within a 1 kb distance from TSS [13].

When looking at the pattern for correlation between DNA methylation and expression, we identified two distinct profiles. We found eQTM associations with a linear distribution between methylation and expression representing a range of continuous values for both DNA methylation and gene expression and, at other loci, associations with a bimodal distribution with no intermediate values for both DNA methylation and gene expression (**Fig 6**). Because DNA methylation is a binary variable as mentioned before, having eQTM associations with bimodal distribution suggest that every interrogated cell within the placenta sample share the same DNA methylation and expression profiles, implying that the association is non cell- or state-specific. On the contrary, eQTM associations with linear distribution suggest a cell- or state-specific mechanism where the continuous signal is explained by the distribution of cells being fully or not methylated among the placenta sample (**S10 Fig**). To assign eQTMs to linear or bimodal distribution we implemented a Bayesian model for bimodal distribution detection. Each gene expression or DNA methylation distribution was analyzed separately and eQTMs were called as linear if both expression and methylation represented a linear distribution and called bimodal if both expression and methylation show a bimodal distribution profile (see Methods).

**Fig 6.**
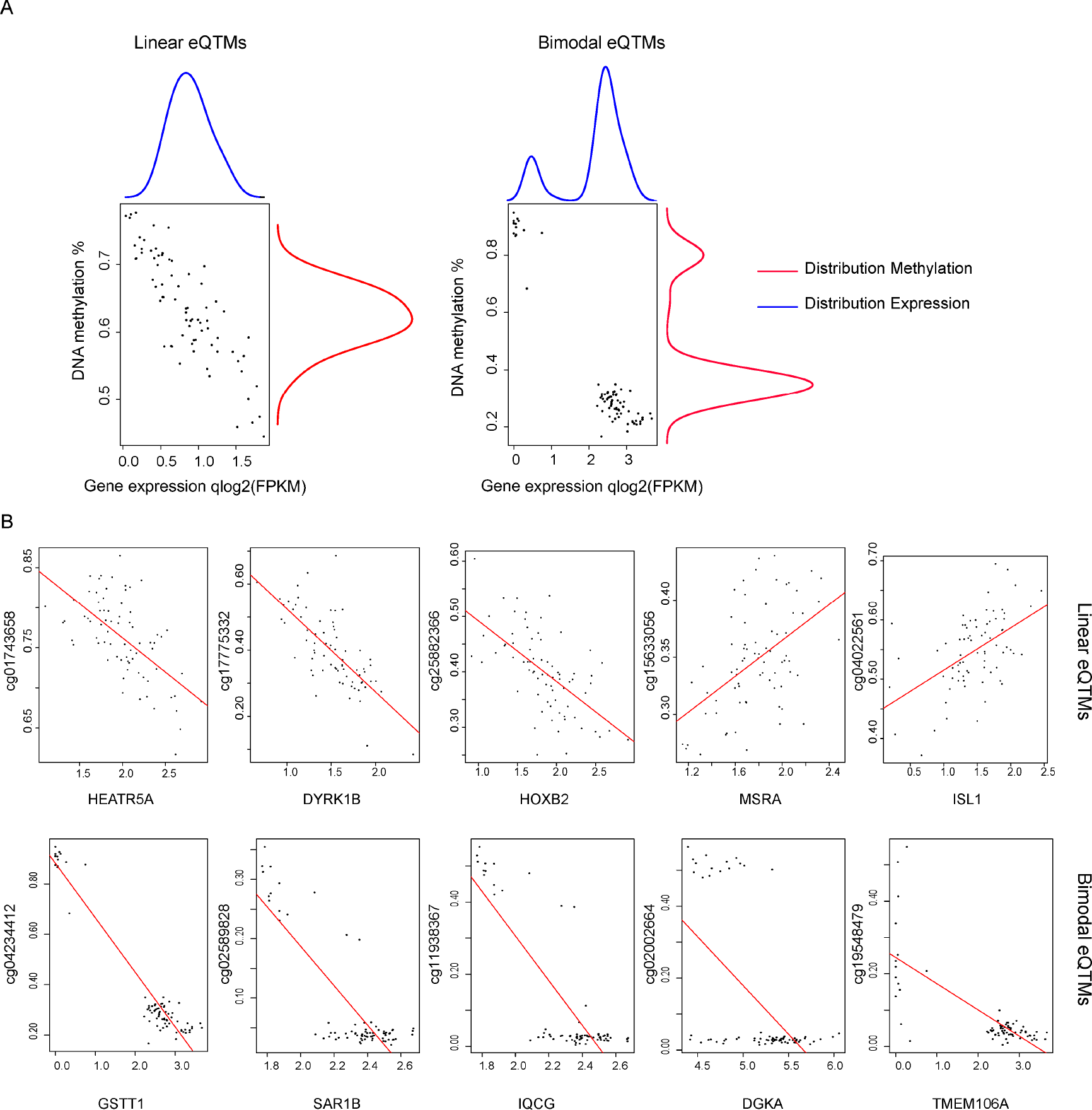
Linear and Bimodal correlations among eQTMs. Representation of linear and bimodal distributions for DNA methylation and expression (A). Scatter plot representing the correlation between DNA methylation and expression among the samples for selected top candidates associations (B). The red line represents the linear regression for each association.

Using our model, we classified 118 associations as bimodal including 29 unique genes and 108 unique CpG sites, and 1,201 as linear which represent 877 unique genes and 1,140 unique CpG sites (**Fig 6**, **S22 Table**). The greater occurrence of linear correlations can be attributed to the high cellular heterogeneity of the placenta. Both linear and bimodal associations were enriched for proximal interactions, with linear showing a more spread distribution (**Fig 7A**). Interestingly, linear and bimodal distributed associations were not found enriched for the same genomic context, linear distributed eQTMs were enriched for feature 2 and 3, our candidate active and poised enhancer and bimodally distributed eQTMs where enriched for feature 1 our candidate promoter (**Fig 7B**).

**Fig 7.**
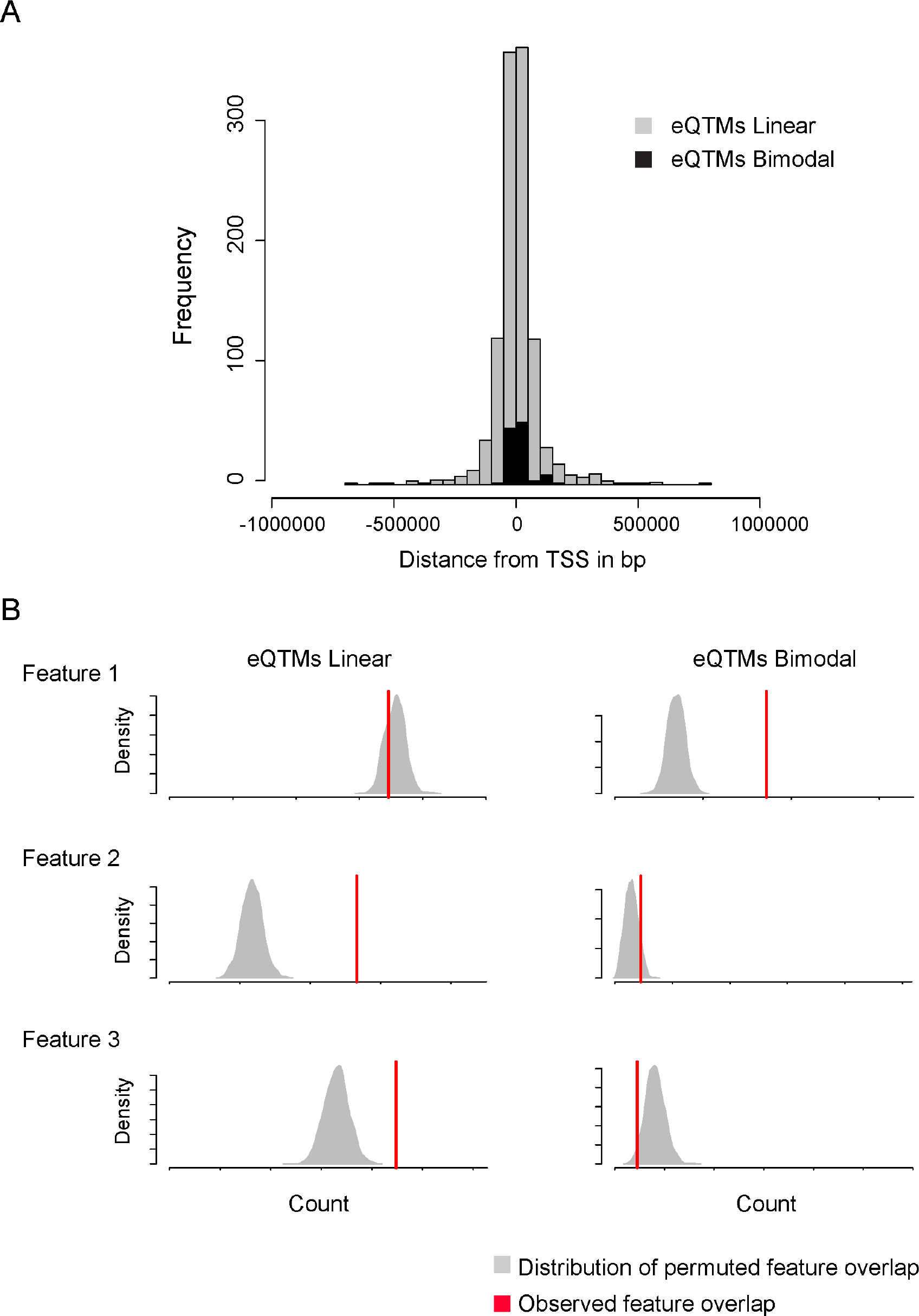
Differences in genomic distribution between linear and bimodal eQTMs. The histogram represents the density of eQTM associations in function of the distance from transcription start site (TSS) considering linear and bimodal correlation separately (A). The density plots represent the distribution of overlaps between random sampling and the different features as defined by permutations tests whereas the red line illustrates the overlap value between candidate associations and features (B).

Looking at transcription factor binding sites across bimodal and linear eQTM associations, we identified a specific pattern where bimodal but not linear associations were globally enriched in transcription factor binding sites (**S21 Table**). Interestingly, the transcription factor previously identified as part of the bHLH family (MAX, MAZ, MXI1, MYC, and SIN3aK20) where highly enriched for bimodally distributed associations while depleted for linear associations (**S11 Fig**). This suggests that distribution not only reflect cell heterogeneity but also provide information on the different factors involved in DNA methylation regulation of expression. These observations, if not sufficient to propose a definitive mechanism of action, will definitely contribute as key resources to further analysis when linking DNA methylation alteration to phenotypes by offering a curated list of gene sensitive to methylation alterations specific to the placenta.

## Discussion

The results presented here highlight the importance of genetic-epigenetic interactions in human placentas and provide a resource to further characterize the dynamic of these interactions in the context of human diseases. Indeed, because GWAS and EWAS analyses are statistical approaches that only report associations between genetic variants and interrogated phenotypes, an approach in which GWAS data are combined with reference datasets detailing associations between genetic variants and gene expression or CpG methylation can help to pinpoint genes and regulatory DNA sequence variants linked to disease susceptibility. By providing a curated list of eQTL, mQTL, and eQTM associations, with careful consideration of potential false-positive and false-negative findings, we have started to address these issues as they pertain to human placentas. Our data represent an initial basis for understanding how genetic variation in human placentas influences epigenetics marks and expression and allows the identification of genes and CpG sites with greater sensitivity to genetic variants, which should be considered when interpreting disease-focused studies such as GWAS and EWAS. Indeed, a significant amount of research today is built on the dichotomy between treated versus non-treated or disease versus non-diseased with less consideration toward understanding what constitutes normal or healthy. Thus, our ability to more fully define a ’normal’ molecular phenotype will have a notable impact on the interpretation and significance of such studies. Lastly, a better understanding of the interplay between the multiple functional layers at a systems level under normal conditions is needed to fully benefit from the ever-increasing amount of multi–omic data.

Epigenome-wide association studies, primarily focusing on DNA methylation, have been used for examining the impact of environmental exposures and early biomarker discovery [13]. The increasing number of studies and challenges in this area have been reviewed [65]. Among these challenges is the ability to adequately correlate DNA methylation and expression profiles to identify reliable and causal associations in a tissue-specific context. Here, we provide a list of candidate genes and CpG sites with high genetic-epigenetic correlations in human placentas, revealing some general principles of these interactions. Associations found in promoter regions represent a categorical/bimodal signal with enrichment for transcription factor binding sites, while enhancer interactions represent a linear additive profile with no significant enrichment for transcription factors. These differences can be related to cell heterogeneity. Indeed, promoter regions show a more constitutive pattern across cell types while enhancers are more likely to be cell type specific. Therefore, enhancer regions will present a cell specific signal depicted as a continuous range of values for gene expression and DNA methylation which transcribe as a linear distribution when looking at methylation-expression associations. Cell subpopulation effects are now recognized as a major source of variability and support the recommendation for single cell type analysis in EWAS [4]. The placenta is a highly heterogeneous tissue and demonstrates differences in gene expression and DNA methylation, which may be due to changes in the proportional distribution of different cell types (for example, as a result of infection, inflammation or other conditions) that can later confound methylation-disease associations [66]. Heterogeneity may be corrected using statistical deconvolution techniques [67, 68] where a subset of targets is used to represent distinct DNA methylation or gene expression profiles for each cell type and assess subpopulation distribution. The accuracy of these approaches is highly dependent on the availability of reference epigenomes and transcriptomes and the ability to select representative targets for the tissue considered. Single cell data for the placenta are not available. However, genes and CpG sites with linear distribution can be further used to assess and control for cell subtype population in human placenta.

There are several strengths to our study, the most important of which is the comprehensive nature of our analysis. By sampling more than 300 individual placentas from an ethnically diverse and representative population, we present here the most extensive placenta-specific genome-wide analysis published to date. Based on our heterogeneous population we believe that our findings will be applicable to populations with diverse backgrounds including minorities that have been so far underrepresented in both GWAS and EWAS studies. Another key factor of our study is the resolution of our approach. We combine from the same cohort genome-wide transcriptome, epigenome and genetic variation information which allows for improved understanding of the intricacy of expression regulation mechanisms in an understudied tissue by overcoming the limitations inherent to each individual dataset when considered separately. Finally, the ability to extract meaningful information is highly dependent on the quality of the analytical approach. Thus, we describe here stringent methods to incorporate different datasets and extract robust correlations. We also emphasize the limitations intrinsic to the different techniques, which will help to assure reproducibility.

A major gap in the field has been the lack of placenta-specific annotations. This situation also makes our findings valuable as they will promote the creation of a placenta-specific data repository. We believe that the massive amount of high-quality data generated will serve as a reference and benefit to further –omics placenta specific studies and contribute to a new perspective in term of disease susceptibility and placenta development.

## Methods

### Study participants

Subjects and samples were identified from the *Eunice Kennedy Shriver* National Institute of Child Health and Human Development (NICHD) Fetal Growth Study, which was a prospective, multicentered, observational study including 3,000 pregnant women [69]. Healthy, non-obese, low risk pregnant women across four race/ethnicity groups, who conceived spontaneously and had no obvious risk factors for fetal growth restriction or overgrowth were eligible for inclusion in the study. Specifically, self-identified non-Hispanic white, African-American, Hispanic, and Asian/Pacific Islander women with a singleton pregnancy less than 13 weeks and 6 days of gestation were enrolled. All women were between 18 and 40 years old with BMI between 19.0 to 29.9kg/m^2^ with no confirmed or suspected fetal congenital structural or chromosomal anomalies. Multiple exclusions were identified to assure a low risk population and included but was not limited to, cigarette smoking in the past six months, use of illicit drugs in the past year, consumption of at least 1 alcoholic drink per day, chronic hypertension, diabetes mellitus, HIV or AIDS, and history of gestational diabetes in a prior pregnancy. To assure correct dating, all pregnancies had first trimester ultrasound screening consistent with gestational age. Placentas were collected from pregnancies with a predicted birthweight by ultrasound between the 10^th^ and the 90^th^ percentile (**S23 Table**).

### Sample collection

Placental samples were obtained and processed within one hour of delivery by trained research personnel. Placental parenchymal biopsies measuring 0.5 cm × 0.5 cm × 0.5 cm were taken from the fetal side of the placenta just beneath the fetal membranes and were placed in RNALater and frozen for molecular analysis.

After delivery, neonatal anthropometric measurements were taken. These included: birth weight, head circumference. Information on maternal weight gain, antenatal and intrapartum complications, and neonatal outcomes were extracted from the medical record.

### Genome-wide gene expression

RNA from 80 placenta biopsies (42 males and 38 females) was isolated using TRIZOL reagent (Invitrogen, MA, USA). Poly-A pull-down was used to enrich for mRNAs, and libraries were prepared using the Illumina TruSeq RNA kit. Libraries were pooled and sequenced on an Illumina HiSeq2000 machine with 100 bp paired-end reads. RTA (Illumina, San Diego, CA, USA) was used for base calling and bcl2fastq (version 1.8.4) for converting BCL to FASTQ format, coupled with adaptor trimming. The reads were mapped to the human reference genome (NCBI/build37.2) using Tophat (version 2.0.4) with 4 mismatches (--read-mismatches = 4) and 10 maximum multiple hits (--max-multihits = 10). The relative expression level of genes was estimated by FPKM (Fragments Per Kilobase of transcript per Million mapped reads) using cufflinks (version 2.0.2) with default settings. FPKM values were used in log2-transformed scale after quantile normalization.

### Genome-wide DNA methylation

Genomic DNA from 303 placental biopsies (151 males and 152 females), 500 ng, was used as per the manufacturer’s instructions for HumanMethylation 450 Beadchips (Illumina), with all assays performed at the Roswell Park Cancer Institute (RPCI) Genomics Shared Resource. Data were processed using Genome Studio, which calculates the fractional methylation (AVG_Beta) at each queried CpG, after background correction, normalization to internal control probes, and quantile normalization. All probes mapping to the X or Y chromosome were removed. As recommended by Illumina, AVG_Beta values with a detection p-value>0.05 were excluded from the analysis and replaced by missing values. Probes that queried CpGs directly overlapping the positions of known common DNA variants as reported in 1000 Genomes Project Phase 3 (allele frequency >1%) and within a 20 base pair window upstream of the interrogated CpG were excluded since these CpGs may be abolished by the SNP itself or the binding of the probe itself could be altered [34]. Additionally, the accuracy of the Illumina 450K can be affected by cross-reactive probes that were also excluded from our analysis [35].

### Genome-wide SNP genotyping

The DNA samples (n=303, 151 males, and 152 females) were genotyped on Illumina HumanOmni2.5 Beadchips, followed by initial data processing using Genome Studio. SNPs were annotated using dbSNP138. Quality control exclusionary measures for subjects were: genotype call rates <95%, marked departure from Hardy-Weinberg equilibrium (p<0.001) and low minor allele frequencies <5% after QC filtering genotype was conserved for 1,374,581 SNPs.

### Association analyses and FDR control

To test for association between genotype and expression (each SNP vs each transcript, eQTL) or genotype and DNA methylation (each SNP vs each methylation site, mQTL) we used a linear regression model implemented in QTLtools [70]. For the association between expression and DNA methylation (each transcript vs each methylation site, eQTM) we used a linear regression model (LINEAR) implemented in Matrix eQTL [71]. For eQTL and eQTM, the linear regression model was corrected using principal component analysis (PCA) and PEER normalization (K=10) based on expression data [72]. For PCA analysis, only the principal components (PC) significantly contributing to variability were added into the model. In our case, PC1 and PC2 were included as contributing for up to 32% of variability (**S2 Fig**). For PEER analysis, covariate contribution was estimated using Bayesian approaches to infer hidden determinants and their effects from gene expression profiles by using factor analysis methods [72]. For each association, FDR was estimated by permutations a 1,000 times after correction for multiple comparisons using Benjamini & Hochberg [73] approach implemented in R package (p.adjust). The same random indexes were applied to the PEER factors and covariates. For eQTLs and mQTLs, permutation is implemented in the QTLtools package. For expression-methylation associations (eQTMs), we permuted ratio for each transcript with significant association (n=14,353) 1,000 times. For eQTL analysis, the correlation was run between expression and genotype across 80 samples (42 male, 38 female), for mQTL analysis association between methylation and genotype was calculated across 303 samples (151 male, 152 female) and for eQTM analysis through 74 samples (38 male, 36 female).

Associations reaching the significance threshold were retained to further analysis if having at least a shared genotype in 5% or more individuals (AA, AB, BB, where A is the reference and B the alternate allele) and regression slope >5%. Associations were further prioritized based on their regression slope and distance between a genetic variant and associated gene or CpG to encompass the linearity and the magnitude of the progression between the different genotype. Briefly, we generated a score for each association following this equation [score = (1-P.value)*abs(regression slope)/log(abs(distance)+2)].

### Determination of “at-risk” probes

“At-risk” probes were defined following 3 distinct criteria (1) the presence of common SNPs in a 20 bp window from the 3’ end of Illumina probe (SNP-probe) [34], (2) previous identification as cross-reactive probes [35] and (3) based on a specific methylation profile using Gap-Hunter algorithm (gap-probe) [36]. Criteria 1 and 2 were sufficient to exclude the probe from further analysis while criteria 3 only resulted in a flagging of the identified probes. We first identified all CpG-SNPs interrogated by the Illumina 450k assay based on 1000 Genomes Project Phase 3 dataset using an allele frequency >1%. We then used the same reference to identify probe with known genetic variants within the 20 bp upstream of the interrogated CpGs including the single base extension for probe of type I. We decided to consider 20 bp because of the absence of consensus for exclusion and because we did not detect a differential enrichment from 20 bp to the 5’ end of the probe (**S12 Fig**). This list of affected probes was further pruned down by excluding probes containing a genetic variant with reference and alternative alleles involving a C or a T which will not affect the binding of the probe as we are considering bisulfite-treated reads. The list of cross-reactive probes is publicly available [35]. We used the function “gaphunter” from the minfi package (v1.18.4) to identify probes with a gap in a beta signal. Such probe signal has a tendency to be driven by an underlying SNP or other genetic variants. Beta values were provided as input and the function outcomes a data frame listing for each identified gap signal, the number of groups and the size of each group. Only gap signals not driven by outlier (cutoff=1%) were considered. A complete list of SNP-probe, cross-reactive probe as well as gap-probe is available as S7 Table. Table S7 also includes the list of association involving a gap-probes. Finally, the non filtered list of mQTLs with annotated probes (SNP/cross-reactive/Gap-probes) is available at S13 Table.

### Enrichment analysis

For an observed overlap X between our candidates and a list of intervals of interest Y of size n from a total population of assayed sites Z, we compared the observed value of X with the distribution of the same overlap n under the null hypothesis using permutation tests. To obtain the null distribution, we employed a random sampling approach where we randomly sampled 1,000 times from the population of all assayed sites (Z). Thus, with Y, n, and an observed overlap with a genomic interval, X, we sampled n loci from Z and found the overlap of the random sample with the interval, Y_k_, for k in 1, 2, …, 1000 (that is 1000 random samples). Next, we compared X with the distribution of simulated overlaps, Y_1, 2, …, 1000_. If the resulting null distribution, Y_1, 2, …, 1000_, contains the observed overlap, X, then we can conclude that there is no significant enrichment. Conversely, when the null distribution, Y_1, 2, …, 1000_, excludes the observed overlap, X, then we can conclude that there is significant enrichment beyond that of random chance. The significance is further assigned as follow: the number of time when Y_k_ > X divided by k.

### π1 statistics

The π1 statistic was used to quantify overlap between eQTL associations found in placenta and the tissue cataloged by the GTEx consortium. The non–model-based pairwise analysis method identifies significant SNP-gene pairs in a first tissue, and then uses the distribution of the *P* values for these pairs in the second tissue to estimate π_1_, the proportion of non-null associations in the second tissue. It provides an estimate of the fraction of true positive eQTLs, π1=1-π0, where π0=estimated fraction of null eQTLs, estimated from the full distribution of p-values (Storey and Tibshirani q-value approach [74]). The π1 statistic considers also sub-threshold placenta eQTL p-values below the FDR<5% cutoff.

### Transcription factor binding affinity estimate

Using the 1000 Genomes Project Phase 3 dataset, we identified all genetic variants within 2 kb windows centered on the gene transcription start site (eQTL) or CpG (mQTL) using a minor allele frequency >1%. The transcription factor dataset was from ENCODE ChIP-seq experiments, together with DNA bindings motifs identified within these regions as displayed by the ENCODE Factorbook repository [75]. We first compare transcription factor binding affinity between the reference and alternative allele using the FIMO tools in the MEME suite [48]. FIMO is a software tool for scanning DNA or protein sequences with motifs described as position-specific scoring matrices. The program computes a log-likelihood ratio score for each position in a given sequence database, uses established dynamic programming methods to convert this score to a *P*-value and then applies false discovery rate analysis to estimate a *q*-value for each position in the given sequence. FIMO output a list of significant binding site for both reference and alternative allele. However, FIMO only provides with qualitative information and does not allow for significant quantification of the difference in binding affinity between the two alleles. Therefore, we used atSNP [49] to quantify the difference between the reference and alternative allele for the binding site identified by FIMO. It uses ENCODE motifs and JASPAR motifs to evaluate the regulatory potential of the SNPs. It outputs for each SNP the significance of the match to each position specific matrix with both the reference and the alternative allele and also the significance of the change in these match scores. atSNP relies on sampling algorithm with a first-order Markov model for the background nucleotide sequences to test the significance of affinity score and SNP-driven changes in these scores. atSNP is an R package.

### Pathways analysis

For pathways analysis, we used the Bioconductor package GOseq [29] combined with the KEGG annotation database [76]. GOseq is based on gene ontology analysis but allows for correction such as gene length that has been shown to bias gene set enrichment analysis. For eQTL associations, we control for gene length as implemented in the reference manual and for mQTL associations, we generated a list of the number of probes per genes using Illumina 450K and Refseq annotations. This list was then used as input to control for gene overrepresentation bias.

### Genome annotation

Placenta specific genome annotation has been generated using publicly available data from the Roadmap in Epigenomics project. Chromatin immunoprecipitation followed by massively parallel sequencing (ChIP-seq) was obtained from fetal placenta primary tissue from several donors (**S16 Table**). Annotation involved processing the raw data for chromatin accessibility (DNase hypersensitivity) along with ChIP-seq data for six histone modifications (H3K4me1, H3K27me3, H3K27ac, H3K4me3, H3K36me3 and H3K9me3), followed by the use of the ChromHMM algorithm [77] to predict seven features as previously described. ChromHMM is based on a multivariate hidden Markov model that models the observed combination of chromatin marks using a product of independent Bernoulli random variables, which enables robust learning of complex patterns of many chromatin modifications. ChromHMM outputs both the learned chromatin-state model parameters and the chromatin-state assignments for each genomic position. Based on enrichment and genomic localization, feature 1 was annotated as candidate promoter (H3K4me3), feature 2 as candidate active enhancer (H3K4me1 and H3k27ac), feature 3 and 4 as candidate poised enhancer (H3K4me1 and H3k27me3) and feature 7 as transcribed sequences (H3K36me3). Finally, features 5 and 6 did not show enough specific enrichment to be annotated (**S7 Fig**).

### Targeted bisulfite sequencing

Targeted bis-seq was performed for the validation of 3 mQTL regions and to assess the effect of common SNPs in 4 Illumina probes as described in Do *et al.* [13]. Briefly, the primers (**S24 Table**) were designed using MethPrimer. Bisulfite-converted DNA was amplified by PCR, followed by Nextgen sequencing (Illumina MiSeq). The PCR and library preparation were performed using Fluidigm Access Array system. PCRs were performed in triplicate and pooled to ensure sequence complexity. ASM was assessed when the coverage was at least 100 DNA fragments. While the absolute differences between methylation of the two alleles are not exaggerated by ultra deep sequencing, the p-values tend to zero as the number of reads increases. Therefore, the Wilcoxon non parametric test was performed using bootstrapping (1000 random samplings, 20 reads per allele) to minimize an artificially low p-value due to ultra deep coverage. The significance of the allele asymmetry was defined by p<0.05 and an absolute methylation difference between allele >20%. For the graphical representations of the bis-seq, one representative random sampling of the reads was shown per allele. Samplings and bootstrapping were performed using R.

### Assessment of the association’s distribution

First, we assessed unimodal and bimodal distribution for expression and methylation data. For expression, a Bayesian model of normal distribution was used to estimate the parameters of the normal distribution fitting the data. The likelihood function is given by a normal distribution:

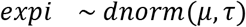

Here, *expi* is the expression value for the *i*^*th*^ sample. *μ* denotes the mean of the normal distribution, *τ* is the precision of the normal distribution. Standard deviation 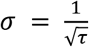.

The prior for the parameter *μ* and *τ* are described by two non-informative priors as:

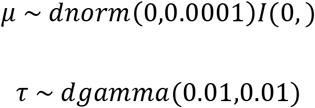

Here, the prior for the *μ* is distributed on the right side of a normal distribution with mean 0 and standard deviation 100. The prior for the precision is described as a gamma distribution with mean 1 and standard deviation 10. To assess the bimodal distribution of expression data, a mixture of two normal distributions model was used to estimate the means, standard deviation and the mass of the bimodal distribution.

The likelihood function can be written as:

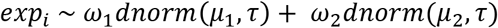

Here, *ω*_1_ and *ω*_2_ represent the mass of each mixture. *μ*_1_ and *μ*_2_ are the means of each mixture. We assumed that the two mixtures have similar variance, *τ*.

The prior for the mass *ω*_1_ is described as:

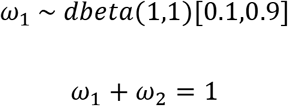

Here, the prior of the mass for the first mixture follows a uniform beta distribution ranging from 0.1 to 0.9, which means the proportion for any mixture has to be larger than 10%. And the two mixtures add up to 1.

The prior for the means *μ*_1_ and *μ*_2_ are described the same way as the mean in a unimodal model.

Because the methylation value for CpG site ranging from 0 to 1, we used a beta distribution to describe the unimodal distribution of the methylation data instead of the normal distribution.

Likelihood:

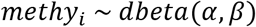

Here, *methy*_i_ is the methylation value for the *i*^*th*^ sample.

The parameters of the beta distribution *α* and *β* are determined by *μ*, the mean of the methylation data, and *κ*, the variation of the methylation data, as:

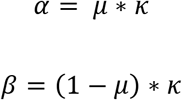

The priors for *μ* and *κ* are modeled as:

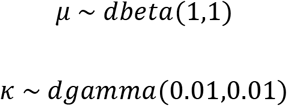

Here, *μ* follows a non-informative uniform beta distribution, and *κ* follows a gamma distribution.

For bimodal distribution of methylation data, a mixture of two beta distributions was used.

The likelihood:

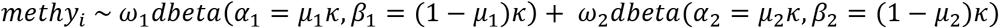

Here, *ω*_1_ and *ω*_2_ represent the mass of each mixture. *μ*_1_ and *μ*_2_ are the means of each mixture. We assume that the two mixtures have similar variance, *κ*.

The prior for the mass *ω*_1_ is described as:

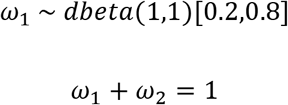

Here, the prior of the mass for the first mixture follows a uniform beta distribution ranging from 0.2 to 0.8, which means the proportion for any mixture has to be larger than 20%. And the two mixtures add up to 1.

The prior for the means *μ*_1_ and *μ*_2_ are described the same way as described in unimodal methylation data.

Having both the model and data, we used the Gibbs Sampler JAGS (Martyn Plummer 2003) to sample the posterior distribution through the “rjags” package (Martyn Plummer 2016). Finally, we compared the goodness of fit of a unimodal and bimodal model for the data by Bayesian Information Criteria (BIC). The unimodality or bimodality of the data is determined by the model with better BIC. The mean and standard deviation of the data is estimated from the posterior distribution sampled by JAGS.

## Acknowledgments

SNP genotyping and methylation Beadchip assays were performed by the Genomics Shared Resource of Roswell Park Cancer Institute, supported by National Cancer Institute (NCI) grant P30CA016056.

## Supporting information

**S1 Fig. Cohort Description.** Heatmap representing gene expression correlation across samples for the full dataset (ALL) and highly variable genes defined using MAD (High MAD) (A). Heatmap representing DNA methylation correlation across samples for the full dataset (ALL) and highly variable genes defined using MAD (High MAD) (B). Genetic background analysis using principal component. Each plot represents principal component 1 versus principal component 2 when considering the 1000 genomes samples reference dataset (1000 Genomes Project samples), our placenta cohort (Placenta Samples) and the overlap between the two. Populations are color coded in the reference dataset and black squares represent samples from the placenta cohort (C).

**S2 Fig. Identification of cofounders.** The principal component analysis was run to identify cofounders within the gene expression and DNA methylation datasets. Histogram representing the proportion of variance explains for each principal component from 1 to 10 (A). Association between principal component and the known factor was assessed using linear regression. Heatmap representing the level of significance for the association between a principal component and each factor for gene expression dataset (B) and DNA methylation dataset (C).

**S3 Fig. Representative association for eQTLs and mQTLs.** Boxplot representing the profile of expression for each genotype for selected top candidate eQTLs (A). Boxplot representing the profile of methylation for each genotype for selected top candidate mQTLs (B).

**S4 Fig. The overlap between placenta specific association and GTEx dataset.** Histogram representing the enrichment for cis-eQTLs from the placenta in transformed fibroblasts from GTEx (A) and in brain cerebellar hemisphere (B). The π1 value represents similarity between tissue ranging from 0 (least similar) to 1 (most similar).

**S5 Fig. mQTL validation.** Targeted bis-seq data showing Hap-ASM in *LOC01848* region. The bis-seq amplicon covers the mQTL index CpG (cg15548566), as well as contiguous CpGs. This region overlaps with the common SNP, rs13360436 which dictates methylation level with the alternate allele (allele B) being significantly hypermethylated compared to the reference allele (allele A), suggesting the presence of hap-ASM in 11 out of 19 heterozygous samples. The low methylated allele is significantly biased toward allele A (p=4×10^−06^, using binomial test) which ruled out imprinting. In this region, rs112724034 has been associated with Alzheimer’s disease (AD) [1]. ∆Meth (difference in the percentage of methylation between alleles in heterozygous samples) and Wilcoxon p-values are from bootstrapping.

**S6 Fig. Example of false-negative in our stringent mQTL set.**Bis-seq showing hap-ASM in *CLDN23* region. cg24900164 was excluded from our main analysis since the probe maps a common non-CT SNP located 7 bp from the index CpG and was therefore not included in our stringent mQTL list. However, targeted bis-seq identified hap-ASM dictated by rs13254997 in 9 out of 20 heterozygous samples (7 positives and 2 negative hap-ASM). In addition, the net methylation estimates from Illumina 450K BeadChips arrays and bis-seq were similar suggesting that the SNP did not affect the probe hybridization. These findings suggest the presence of genuine mQTL at this locus.

**S7 Fig. ChromHMM annotations.** ChromHMM algorithm was used to define genomic annotations based on ChIP-seq tracks available for the placenta. Heatmap representing the enrichment for the different ChIP-seq mark in each feature (A). Heatmap representing the enrichment for previously defined genomic annotation in each feature (B). Density plot representing the enrichment for each feature in function of the distance from the transcription starting site (TSS) (C).

**S8 Fig. Enrichment for proximal associations.** Histogram representing the distribution of associations from the transcription start site (TSS) for eQTL (A) and from the CpG site for mQTL (B).

**S9 Fig. Distribution of difference in binding affinity.** Representative density plot of the difference in binding affinity between the reference and alternative allele for eQTL (A) and mQTL (B) as defined by FIMO. Null differences and binding not reaching a p.value <0.0001 were excluded prior to analysis.

**S10 Fig. Schematic representation of the subpopulation effect on the correlation between DNA methylation and gene expression.** Scenario representing a correlation between CpG and gene which is not cell specific as DNA methylation and gene expression profiles are similar across the different cells of each sample. CpG is either fully or not methylated and expression is high or low. This scenario will be called as bimodal distribution (A). Scenario representing an absence of correlation between DNA methylation and gene expression (B). Scenario representing a correlation between CpG and gene which is cell specific as DNA methylation and gene expression profiles are variable across cell within each sample. This profile will be called a linear distribution (C).

**S11 Fig. Transcription factor binding sites enrichment for eQTMs.** The density plots represent the distribution of overlaps between random sampling and the selected transcription factors binding site as defined by the ENCODE Factorbook repository when the red line illustrates the overlap value between candidate associations and the corresponding transcription factor binding site for linear (A) and bimodal (B) distributions.

**S12 Fig. mQTL association within Illumina 450k probes.** Scatter plot representing the enrichment for significant associations in function of distance from 3’end of Illumina probe. The blue dash line represents the cutoff use during our study. This cutoff was defined based on previous observations and on the decrease, enrichment observed in our analysis.

**S1 Table. Correlation of expressions between placenta and Tissue from the GTEx consortium.**

**S2 Table. Summary of genes with high variability across the placenta.**

**S3 Table. Summary of CpGs with high variability across the placenta.**

**S4 Table. Summary of pathways associated with gene showing high variability.** This table relates the significant outcome of the gene set enrichment analysis using GOseq and KEGG database for genes and CpG-associated genes classified as high variable across placenta using MAD.

**S5 Table. Summary of eQTL associations.**

**S6 Table. Summary of mQTL associations.**

**S7 Table. Excluded probes.** This table recapitulates the list of probes from Illumina 450K design that were excluded in the present analysis, the list of cross-reactive probes as defined by Chen YA, *et al.*, the list of probes that were flagged by Gap-hunter as well as the list of mQTLs and eQTMs that match with these flagged probes.

**S8 Table. Summary of genes associated with eQTL overlapping with the previously published association by Peng *et al.*.**

**S9 Table. mQTL association overlapping with previously identify mQTLs in the Do *et al.* paper.** The overlap between our candidate mQTLs and the one previously identified by Do *et al.* was generated focusing on mQTL associated CpG.

**S10 Table. π1 score across the GTEx tissues.** π1 was calculated for each available sample from the GTEx database as a measure of similarity between these tissues and placenta.

**S11 Table. List of placenta specific associations.**

**S12 Table. mQTLs validation.** The outcome of the validation using a targeted approach for selected mQTL candidate found in our and Do *et al.* studies.

**S13 Table. The complete list of mQTL associations with flagged at “risk” probes.**

**S14 Table. KEGG analysis.** This table relates the significant outcome of the gene set enrichment analysis using GOseq and KEGG database for eQTL and mQTL associations.

**S15 Table. List of genes overlapping between eQTLs and mQTLs.**

**S16 Table. ChromHMM input dataset.** Information regarding placenta specific ChIP-seq tracks used to perform ChromHMM.

**S17 Table. List of the transcription factor with altered binding affinity due to genetic variants associated to eQTLs and mQTLs.** This list represents the outcome of the atSNP algorithm.

**S18 Table. The overlap between placenta specific eQTLs and associations from the GWAS database.** This analysis was centered on overlapping genetic variants between placenta eQTLs and genetic variant previously reported in the GWAS database.

**S19 Table. The overlap between placenta specific mQTLs and associations from the GWAS database.** This analysis was centered on overlapping genetic variants between placenta mQTLs and genetic variant previously reported in the GWAS database.

**S20 Table. Summary of eQTM associations.**

**S21 Table. Transcription binding sites analysis for eQTM associations.** List of transcription factor binding sites showing significant enrichment for eQTM candidates considering bimodal, linear distribution and positive or negative correlation separately.

**S22 Table. Summary of eQTM association based on the correlation distribution profile.**

**S23 Table. Cohort information.**

**S24 Table. List of primers for bis-seq validation.**

Codes used during our analysis are available here: [https://github.com/fabiendelahaye/Placenta_analysis.git]

